# Direct brain recordings identify hippocampal and cortical networks that distinguish successful versus failed episodic memory retrieval

**DOI:** 10.1101/2020.05.27.120097

**Authors:** Ryan Joseph Tan, Michael D. Rugg, Bradley C. Lega

## Abstract

Human data collected using noninvasive imaging techniques have established the importance of parietal regions towards episodic memory retrieval, including the angular gyrus and posterior cingulate cortex. Such regions comprise part of a putative core episodic retrieval network. In free recall, comparisons between contextually appropriate and inappropriate recall events (i.e. prior list intrusions) provide the opportunity to study memory retrieval networks supporting veridical recall, and existing findings predict that differences in electrical activity in these brain regions should be identified according to the accuracy of recall. However, prior iEEG studies, utilizing principally subdural grid electrodes, have not fully characterized brain activity in parietal regions during memory retrieval and have not examined connectivity between core recollection areas and the hippocampus or prefrontal cortex. Here, we employed a data set obtained from 100 human patients implanted with stereo EEG electrodes for seizure mapping purposes as they performed a free recall task. This data set allowed us to separately analyze activity in midline versus lateral parietal brain regions, and in anterior versus posterior hippocampus, to identify areas in which retrieval–related activity predicted the recollection of a correct versus an incorrect memory. With the wide coverage afforded by the stereo EEG approach, we were also able to examine interregional connectivity. Our key findings were that differences in gamma band activity in the angular gyrus, precuneus, posterior temporal cortex, and posterior (more than anterior) hippocampus discriminated accurate versus inaccurate recall as well as active retrieval versus memory search. The left angular gyrus exhibited a significant power decrease preceding list intrusions as well as unique phase-amplitude coupling properties, whereas the prefrontal cortex was unique in exhibiting a power increase during list intrusions. Analysis of connectivity revealed significant hemispheric asymmetry, with relatively sparse left– sided functional connections compared to the right hemisphere. One exception to this finding was elevated connectivity between the prefrontal cortex and left angular gyrus. This finding is interpreted as evidence for the engagement of prefrontal cortex in memory monitoring and mnemonic decision–making.

## 1. Introduction

Numerous non–invasive imaging studies have provided evidence for the importance of parietal lobe locations to the retrieval of episodic memories. Both midline and lateral regions, namely the posterior cingulate cortex and angular gyrus, are critical hubs within a putative core episodic memory retrieval (recollection) network, along with medial prefrontal areas, the parahippocampal cortex, and temporal areas.^26,50,10,19,31^ These studies have demonstrated that activity in this network covaries with the amount of retrieved information and that the content of retrieved memory items can be decoded from activity in this network. Remarkably, parietal regions within this network also exhibit retrieval–related activation that is independent of the specific type of memory item (i.e. spatial information, verbal memories, and autobiographical memories).^1,2,27,42^ Studies using intracranial EEG have shown that autobiographical memory retrieval modulates medial–lateral parietal connectivity.^13^

Based upon the properties of the recollection network determined using noninvasive studies, especially the observation that network regions are sensitive to the fidelity of recollected information,^43^ one would predict that activity within this network would distinguish correct versus contextually inappropriate memory retrieval in episodic memory paradigms. Contrary to this prediction, however, previous analyses of high–frequency activity (HFA defined as gamma band power: 44–100 Hz) during episodic memory retrieval suggest that the parietal cortex exhibits a nonspecific HFA increase and only weakly discriminates between contextually appropriate versus inappropriate recollection^37^ (using list intrusions to operationalize contextually inappropriate recollection). The somewhat surprising implication of these data (in light of these aforementioned models of a recollection network) were that the hippocampus, but not the parietal cortex, represents contextual information in episodic memory. This tension between iEEG observations and models derived principally from noninvasive human data was the central motivation for our analysis, as we believe that the aggregation of electrodes from multiple parietal locations (including Rolandic, opercular, angular, supramarginal, SPL, and midline structures from both hemispheres) attenuated dissociable contributions from ventral versus dorsal and medial versus lateral parietal lobe locations. In addition, no studies have characterized brain *connectivity* during correct versus failed recollection using intracranial EEG. Investigating electrophysiological patterns of fronto–parietal connectivity during episodic memory could test how frontal regions (such as the dorsolateral prefrontal cortex) interact with the recollection network to provide a hypothesized monitoring function during episodic retrieval.^3,18,4^

The features of our dataset, including simultaneous frontal and parietal recording locations, bore the potential to link together these complementary theoretical treatments by identifying specific areas within the recollection network that are potential targets of frontal lobe modulation. Finally, previous investigations have not compared retrieval–related activity between the anterior and posterior hippocampus for correct versus failed recall. Examination of this issue with more precise electrode localization than has been achieved previously can inform theories of hippocampal longitudinal specialization, and build on our prior work suggesting that retrieval–related processes preferentially activate posterior hippocampal locations.^36,35^

With the goal of providing a more precise anatomical characterization of electrophysiological activity during episodic memory retrieval (including connectivity information), we analyzed a data set derived from 100 subjects who performed free recall after implantation of intracranial electrodes via the stereo EEG technique. We utilized prior list intrusions in the free recall paradigm as a behavioral contrast;^23,44,37^ we compared HFA aggregated across subjects within one of nine predefined regions of interest in both hemispheres. The regions were the anterior hippocampus, posterior hippocampus, angular gyrus, supramarginal gyrus, posterior cingulate, precuneus, anterior temporal cortex, posterior lateral temporal cortex, and dorsolateral prefrontal cortex. Thus, we included regions both within and outside of the core episodic recollection network. We tested hypotheses directly motivated by previous human noninvasive experiments, namely the putative importance for accurate memory retrieval of left posterior parietal cortex, midline parietal regions, and the hippocampus. Although we did not have sufficient representation in medial prefrontal regions or parahippocampal cortex to characterize the entire recollection network, we were able to take advantage of the broad coverage afforded by stereo EEG to analyze connectivity among network members including the hippocampus and angular gyrus. We also analyzed anterior temporal and dorsolateral prefrontal regions to provide a contrast with activity from core recollection network regions and look for evidence of modulation of prefrontal connectivity according to retrieval success. We also directly contrasted left versus right angular gyrus activity and network connectivity separately for left and right hemispheres, as well as comparing anterior versus posterior hippocampal activity. In addition, we analyzed phase–amplitude coupling during contextually appropriate versus inappropriate retrieval. We discuss how our data fit with different views of the processes underlying prior list intrusions, including the involvement of prefrontal control mechanisms.^21,41^ We situate our findings within the context of existing theories of episodic memory retrieval^21,23,37^ and hippocampal longitudinal specialization.^36,35^

## 2. Materials and Methods

### 2.1. Participants

100 subjects with medication resistant epilepsy who underwent stereoelectroencephalography surgery with the goal of identifying their ictal onset region(s) participated in the study. Participants came from the University of Texas Southwestern (UTSW) epilepsy surgery program across a time span of 4 years. Subjects had intracranial depth electrodes implanted at locations specified by the neurology team, and electrodes were laterally inserted into the specified regions with robotic assistance.^15^ Final electrode localization in the frontal, temporal, and parietal subregions was determined by post–operative expert neuroradiology review. We obtained written and informed consent from each patient. Protocols were approved by the UTSW Institutional Review Board on Human Subjects Research. In 8 subjects, the language dominant hemisphere was determined to be bilateral or could not be ascertained via preoperative fMRI; the remainder were determined to have left hemisphere dominant language function. Subjects able to complete one full session of free recall with at least 10% correct retrievals were included. Seizure localization in such a large dataset was quite diverse, including frontal, temporal, and occipital onset locations. Electrodes from hemisphere of evident radiographic abnormalities including temporal sclerosis or previous neurosurgery were excluded from analysis. We aggregated electrodes into a set of nine regions, namely the anterior hippocampus, posterior hippocampus, angular gyrus, supramarginal gyrus, posterior cingulate, precuneus, anterior temporal cortex, posterior lateral temporal cortex, and dorsolateral prefrontal cortex. All regions had at least 15 subjects contributing data to each region. Depth electrodes were Ad-tech or PMT depth electrodes 0.8 mm in outer diameter, with form 10 to 14 recording contacts arrayed at 4-5mm intervals along the shaft of the electrode (number of contacts is uniformly 10 with variable spacing for Ad-tech electrodes, but 10-14 with uniform spacing for PMT electrodes depending upon the overall depth of insertion).

**Table 1:** The number of subjects and electrodes included in this study per region.

| Region | No. of Subjects | No. of Electrodes |
| --- | --- | --- |
| AH | 51 | 209 |
| PH | 46 | 148 |
| AG | 33 | 169 |
| SMG | 41 | 206 |
| Prec | 43 | 119 |
| PC | 52 | 108 |
| AT | 49 | 468 |
| PLT | 48 | 293 |
| LPFC | 41 | 337 |

### 2.2. Free Recall Task

Participants performed a verbal free recall task, in which lists of 12 or 15 high–frequency, non–semantically related nouns were chosen from a pool without replacements (http://memory.psych.upenn.edu/Word_Pools)^44,37,46^ and presented on a screen one at a time for 1.6 seconds, each followed by a 1 second blank screen. Each list was followed by a 30 second math distractor period where participants answered arithmetic problems in the form of A + B + C = ??. Participants were explicitly instructed to recall only items from the immediately preceding list and not from other lists. After the distraction period, participants attempted to recall as many words as possible from the list just presented to them in no particular order within a 30 second retrieval period. This procedure was repeated 25 times per session, and each participant completed on average 2 sessions throughout their stay with unique lists. Recall responses were manually scored to identify correct retrievals versus intrusions. Extra list intrusions (items never presented on other lists) were separated from prior list intrusions (items presented on previous lists). In the analyses described below, we included contrasted activity associated with correct recalls and prior list intrusions (PLIs) only.

### 2.3. Retrieval Data Analysis

Intracranial EEG was sampled at 1kHz on a Nihon–Kohden 2100 clinical system under a bipolar montage with the most medial white matter contact on individual electrodes as reference. We applied an offline noise detection algorithm to filter out noise and possible interictal activity.^32^ EEG data were sampled at 500Hz. We analyzed two classes of retrieval events in our analysis: correctly retrieved items and PLIs. In addition, we selected other epochs to represent deliberation activity (see below). Retrieval events had to be preceded by at least 1500 ms in which no other verbalization occurred to be included in the analysis, following previously published methods.^37^ For this analysis, 74% (8063 out of 10969) of these retrieval events met this criterion. For both correct retrieval and PLI events, the analysis epoch comprised the 1000 ms preceding the onset of vocalization. Deliberation events,^37^ comprised 500 ms epochs selected at random from the retrieval period with the constraint that they were preceded and followed by 1500 ms in which no vocalization occurred. For the analysis of these deliberation periods, we precisely followed previously published methods.^5,37^ Each included participant met a threshold of at least 10 correct retrievals and 10 PLIs to be considered in the analysis.

### 2.4. Oscillatory Power Analysis

Oscillatory power at 46 logarithmically spaced frequencies between 2 to 100 Hz was extracted using a Morlet wavelet transformation (wave number = 6) applied to the EEG obtained from all electrodes during pre–retrieval and deliberation epochs (see above). To avoid possible edge effects, a 1000 ms buffer was applied to each side of the signal.^37^ Power values were also extracted 200 ms before word onset during the encoding period and used to normalize the power of the retrieval events relative to the mean and standard deviation of the encoding events. Normalization was carried out within each experimental session, following previously published methods.^37^ Retrieval events of each category (correct retrievals and PLIs) were combined by averaging all event power values within each electrode. For each subject, all possible electrodes within a region of interest were aggregated so that a subject contributed a single data point for each region of interest. Each subject’s data was then averaged across gamma band frequency (44-100 Hz) and time (1000 ms before vocalization). We then aggregated all subjects within a region and directly compared the distributions of gamma activity values for PLIs and correct retrieval events using a pairwise t–test. This gamma frequency range was used to match previously published methods and to limit the number of multiple comparisons, with this band thought to serve as a proxy for neural activation.^37^ To calculate significance values, we performed a within region shuffle procedure to generate a distribution of 1000 p values and compared it to the true distribution to determine the likelihood that a correct retrieval versus PLI difference in a each region occurs by chance. The final p values were corrected for multiple comparisons using the false discovery rate (FDR). This procedure was repeated for the deliberation analysis, for which the number of deliberation events was matched to the number of correct retrievals by randomly selecting 1000 msec epochs separated from vocalization events.^37^ Deliberation events were compared to both correct retrieval and PLI events individually, and the final p values were corrected for multiple comparisons as well.

### 2.5. Phase–Amplitude Coupling

Phase–amplitude coupling was extracted using the modulation index method.^47^ The phase from two low–frequency bands (2–5, 4–9 Hz) and the amplitude from a gamma band range (40– 70 Hz) was extracted using a Hilbert transformation during the correct retrieval and PLI event period.^12,34^ Two theta bands were used because of existing observations that hippocampal PAC occurs in the lower frequency range, while in the cortex it occurs in the higher range.^34^ The phase–amplitude distribution was then constructed by first creating 18 equally spaced phase bins between −*π* and *π* radians. We then separated the amplitude of each sample in accordance with the phase bin of the low–frequency band. We then normalized the amplitudes within each bin by dividing by the total averaged amplitude. Shannon entropy and subsequently the Kullback–Leibler distance were calculated, and the MI was obtained by dividing the Kullback– Leibler distance by the maximum entropy value.^47^ MI values were calculated on a trial-by-trial basis, normalized using a z-transformation, and averaged within electrode. For each subject, all possible electrodes within a region of interest were aggregated so that a subject contributes a single data point for each region of interest. We then aggregated all subjects within a region and MI values were compared using a pairwise t-test. Like the power analysis, we utilized a within region shuffle to compare the true statistic to the distribution of 1000 null statistics generated to establish significance. We adopted a strict threshold of .01 to determine significant PAC. The preferred phase was extracted by taking the mean phase of correctly retrieved events in electrodes that showed a significant MI difference between the two conditions. Specifically, we used the real and imaginary components of the phase distribution as independent variables in a regression equation with the amplitude as the dependent variable. We converted the resulting regression coefficients to obtain a representation, across time, of a single phase value that best predicted power at higher frequencies (40-70 Hz) within each electrode.^34^ After aggregating across subjects, we tested the phase distribution using a Rayleigh test.^40^

### 2.6. Connectivity Analysis

In this analysis, we sought to identify functional connectivity, meaning phase coherence between two regions that is significantly different depending upon whether an individual retrieves a correct item or a PLI. Functional connectivity measurements were extracted using the phase locking value (PLV) method.^33^ Phases were extracted from all trials for each region-region pair. PLV was calculated by calculating a mean resultant vector length (Equation 1) across all trials for both correct retrieval and PLI conditions. A PLV difference between the two conditions was calculated to establish a functional connectivity metric.^46^ Each electrode pair underwent a permutation test of significance by applying the shuffling procedure described previously^46^ to obtain a single phase locking statistic (PLS) distribution for each electrode pair (a functional connectivity measurement). PLS values were normalized using a z-transformation, then averaged across all possible pairwise electrode combinations within a subject so that each subject only contributed one data point for each region-region comparison. Each subject’s data was then average across theta band frequency (3–9 Hz) and time (1000 ms before vocalization). The functional connectivity was averaged across this theta frequency band to follow previously reported studies and limit the number of multiple comparisons.^46^ To test for network-wide significance, we generated 1000 null PLS distributions derived from the shuffled trials to generate chance network statistics. The null distribution underwent every operation as the true PLS distribution. PLS values from the PLI/correct retrieval comparison for a given connection underwent a permutation test and were compared to the null distribution to establish network-level significance. Overall, larger z-PLS value indicated greater functional connectivity in correctly retrieved trials, while a smaller z-PLS value indicated greater functional connectivity in intrusion trials. We adopted a strict threshold of .01 to determine significant connectivity between region-region pairs. Phase lag was calculated by identifying the peak time of the functional connectivity value for each region–region electrode pair, and extracting the mean relative phase across all correctly retrieved events at that time point. Mean phases were then aggregated across electrodes and analyzed using a Rayleigh test. *Note on theta frequency ranges: Based on existing data for PAC, we separated the phase-providing frequency range into two separate bands for this analysis. However, for the connectivity analysis, we used a single broad 3–9 Hz frequency in order to precisely follow previously published methods.*

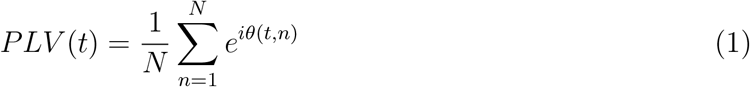

## 3. Results

### 3.1. Comparison of HFA during correct versus incorrect memory retrieval

We investigated the electrophysiology of episodic memory retrieval by comparing gamma (44–100 Hz) power (HFA) preceding correctly recalled items and prior list intrusions in a free recall task. Our dataset (consisting of 100 unique individuals) permitted us to differentiate patterns among parietal lobe locations in both hemispheres. We aggregated electrodes into a set of nine regions, namely the anterior hippocampus (AH), posterior hippocampus (PH), angular gyrus (AG), supramarginal gyrus (SMG), posterior cingulate (PC), precuneus (Prec), anterior temporal cortex (AT), posterior lateral temporal cortex (PLT), and dorsolateral prefrontal cortex (DLPFC). The number of individuals contributing electrodes to each of the regions is included in Figure 1. We analyzed the serial position of correctly recalled items and list intrusions in both encoding and retrieval to distinguish any behavioral effects between the two retrieval classes. We observed a strong PLI primacy effect in both the encoding serial position and the position that the word was retrieved. The proportion of PLIs stemming from the encoding of primacy items significantly exceeded that expected by chance (*p* < 0.001, binomial test), but was not different than the proportion of correct retrievals made from primacy items overall (p = 0.48). 28% of PLI events occurred in the first two output positions (Figure 1E), a proportion significantly less than the proportion of correct retrieval events occurring in the first two output serial positions (*χ*^2^ = 93.64, p< 0.001). However, to look for any confounding effect of serial position, we compared HFA during PLI events for early (first two) versus later serial positions. Across all regions, there was no significant difference (t(445)=–0.0772, *p* < 0.5308). Furthermore, we compared all PLI events to both early and late correct retrieval events. We found both early and late correct retrieval events were significantly different than PLI events (t(982)=3.6899, *p* < 0.0001 and t(982)=3.7070, *p* < 0.0001, respectively).

**Figure 1:**
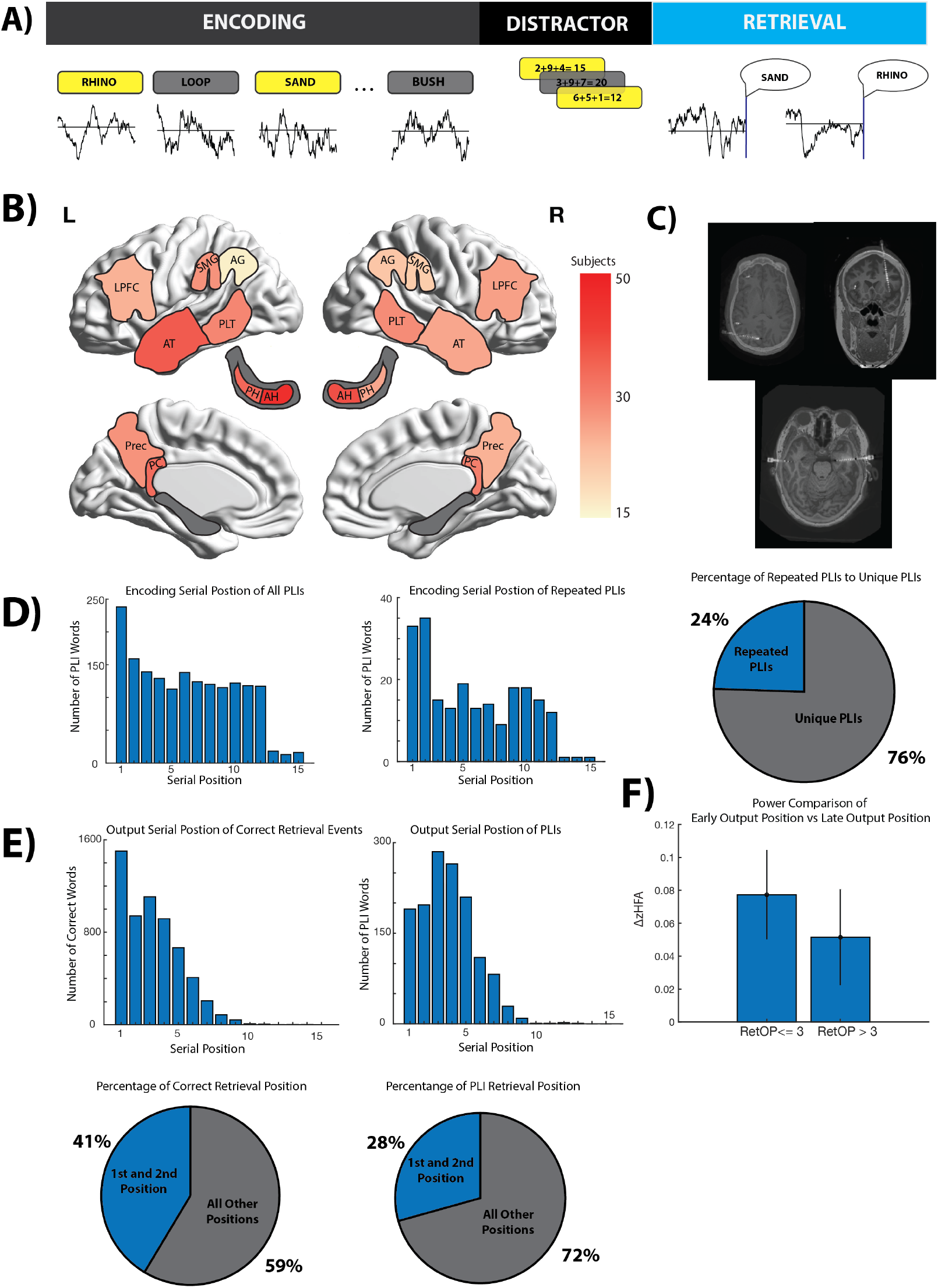
Summary of free recall task and electrode distributions for subjects. (A) Example of a single free recall trial. The time bin from 1000 ms before retrieval vocalization was used for analysis. (B) Brain plot showing the number of subjects per given brain region. Each region had a minimum of 15 subjects included in the analysis. All analyses conducted at the subject level (single data point per subject per region). (C) CT scan showing laterally inserted electrodes from a representative subject. (D) Histogram of PLI words based on encoding serial position and proportion of repeated PLIs retrieved to unique PLIs retrieved. (E) Histogram of output serial position for both event categories and proportion of first and second retrieved items (first two output serial positions) at which correct retrievals and PLIs occurred. (F) Comparison of the delta power of retrieval events with output serial position (OSP) <= 3 and OSP > 3. Error bars are standard error of the mean. This was not significant (*p* < 0.4582).

### HFA differences characterize a memory retrieval network

We sought to define a retrieval network according to two criteria: 1) significantly greater oscillatory power preceding correct retrieval as compared to list intrusions and 2) greater oscillatory power preceding correct retrieval as compared to a deliberation condition in the retrieval phase of the free recall task.^37^ The deliberation periods comprise epochs during which no recalled item was vocalized within 1500 msec, and in prior publications, these have been characterized as periods of ‘memory search.’^37^ All power values were normalized using a period of a blank screen during item encoding following previously published methods.^37^ Results are shown in Figure 2. Regions that exhibited a significant difference in HFA included left and right posterior (but not anterior) hippocampi, right angular gyrus, bilateral precuneus and right posterior lateral temporal cortex. These findings are consistent with previous reports in that we observed a significant HFA difference in the hippocampus,^37^ but here this hippocampal effect appears to be driven by greater retrieval–related differences in the posterior hippocampus. Using the data from those subjects in whom there were both anterior and posterior hippocampal contacts (N= 41, Figure 2B), a pairwise t–test with shuffle procedure indicated that the posterior hippocampal power increase was significantly greater (t(40) = 2.9932, *p* = 0.0018). Power differences were broadly significant across core retrieval network locations,^42^ and, notably, were specific for the angular but not the supramarginal gyrus. We directly compared power during correct retrieval for the angular versus supramarginal gyri, as these regions have previously shown to exhibit different patterns during memory retrieval. We found the angular gyrus power increase to be significantly greater(t(14)=1.9041, *p* = 0.0336). While only the right angular gyrus survived FDR correction for both comparisons (retrieval vs. PLI, retrieval vs. deliberation), the pattern of the effect was similar and direct left versus right angular gyrus comparison was not significant (t(31) = 1.1666, *p* < 0.1261).

**Figure 2:**
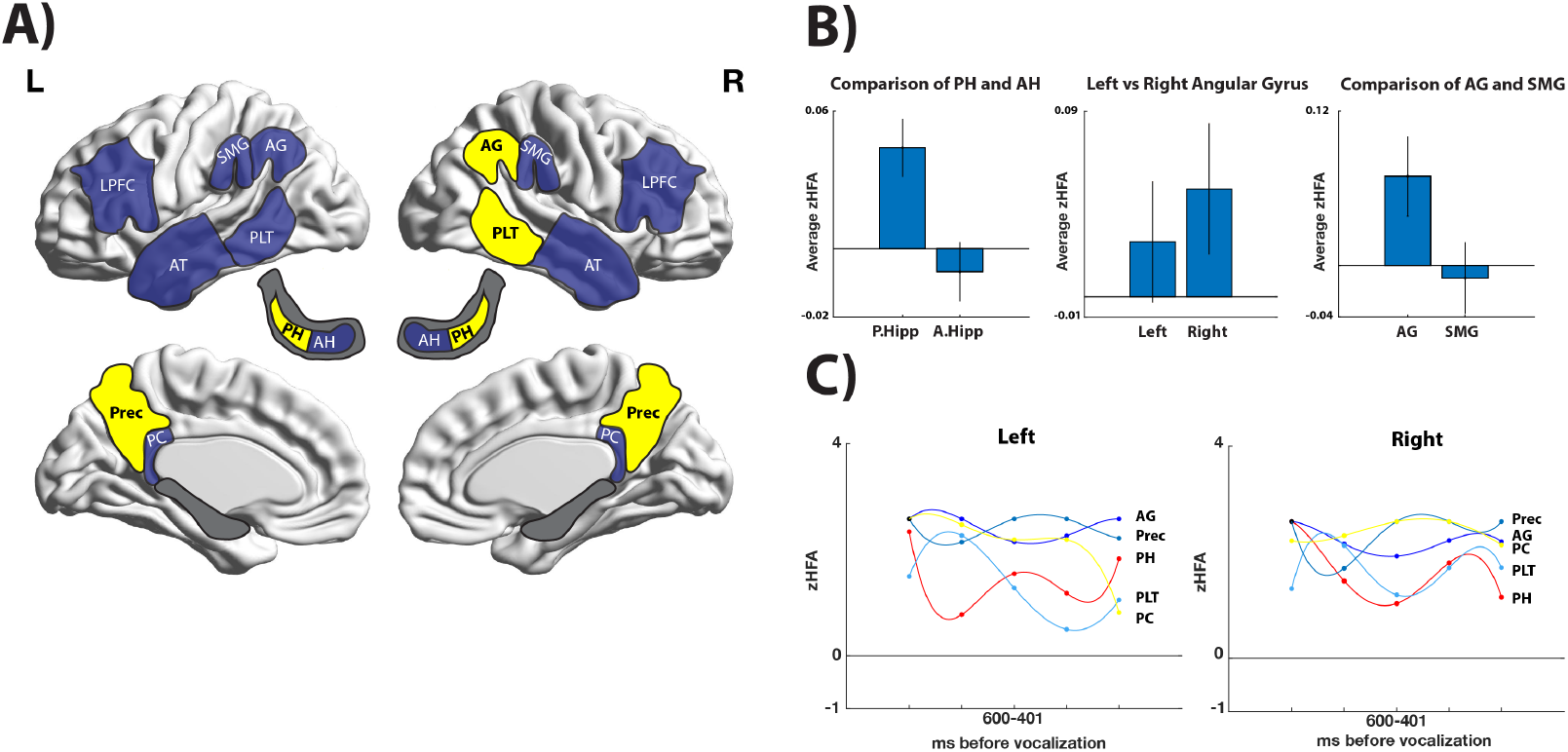
Overview of high frequency activity during correct retrieval. (A) Brain plot showing (in yellow) regions that show a significant power increase during correct retrieval events in compared to both PLI and deliberation events. (B) Individual region comparison plots comparing average normalized power during correct retrieval events. Error bars are standard error of the mean. PH vs AH power as well as AG vs SMG power was significantly different (*p* < 0.0018 and *p* < 0.0336 respectfully). Left vs Right AG power was not significant (*p* < 0.1261).(C) Mean difference of HFA during correct and incorrect memory retrieval sorted in 200ms time bins. A best fit line (4th degree polynomial) was applied to each region to show the temporal dynamics of the HFA differences during retrieval.

We observed a relative decrease in HFA during erroneous retrieval common throughout parietal brain locations as well as the anterior but not posterior hippocampus. We noted a significant HFA power decrease in the left angular gyrus, whereas the opposite pattern was observed in the bilateral prefrontal cortex, in which both successful retrieval and list intrusions exhibited a relative HFA power increase. This is visible in Figure 3, which shows the HFA patterns across the brain for each condition. We tested for differences across all brain regions in these PLI HFA values, revealing a significant effect of brain region across all subjects (F(8,508) = 2.64, p = 0.007) with a significant difference observed between right prefrontal (HFA increase during PLI) and left angular gyrus (HFA decrease during PLI) (t(42) = 3.7858, p = .00024). We plotted the timecourse of PLI versus correct retrieval HFA differences in Figure 2 by separately comparing HFA distributions across subjects after binning recorded power values (200 msec bin width). For the angular gyrus, this revealed a stable pattern across time, from 1000 msec to 200 msec prior to vocalization of retrieved items. In the posterior hippocampus, by contrast, there was a bimodal response, with a peak in PLI/correct differences occurring earlier in time (1000 msec before vocalization) and again immediately preceding vocalization.

**Figure 3:**
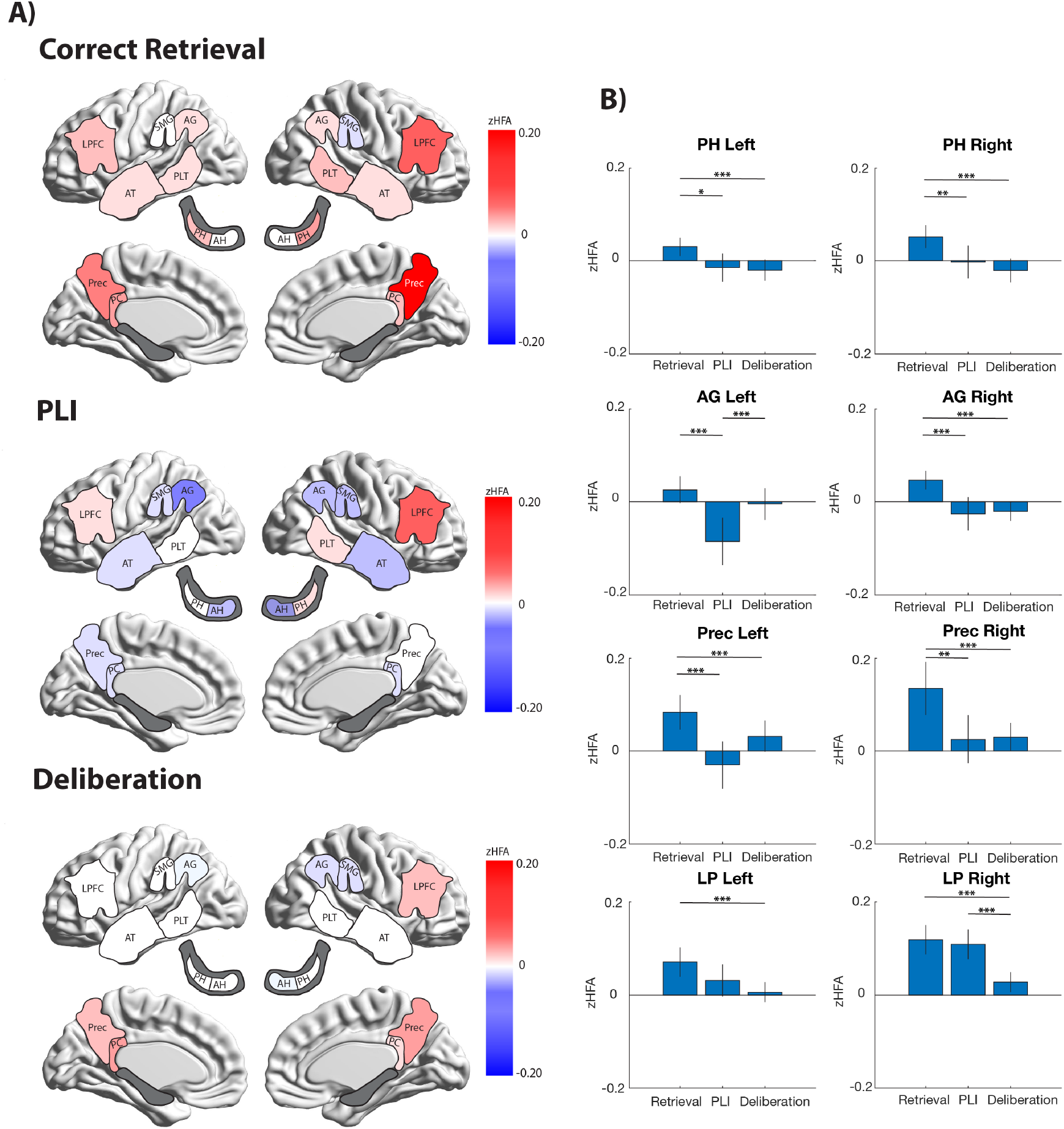
High–frequency activity during correct and incorrect memory retrieval. (A) Brain plots of average zHFA for Correct Retrieval, PLI, and Deliberation condition. (B) Bar plots depict average zHFA in Retrieval, PLI, and Deliberation epochs (corrected *p* < 0.05, t–test with shuffle procedure). Astricks denotes significance (* <.05, ** < 0.01, *** < 0.001)

In contrast to the above pattern, we observed a significant difference between deliberation and PLI activity in the left posterior cingulate region, driven by power increase during the deliberation phase rather than relative suppression of HFA preceding list intrusions (t(32) = 2.5835). This pattern (strong HFA increase during the deliberation period) is relatively unique across the data set, possibly reflecting earlier onset of retrieval related activity in the posterior cingulate cortex rather than an HFA power increase immediately preceding item retrieval. The precuneus exhibited a weaker HFA increase during deliberation, but notably this region in both hemispheres exhibited the strongest HFA power increases for correct retrieval events (F(8,508) = 3.94, p = 0.002). This is visible in Figure 3.

### 3.2. Phase–Amplitude Coupling

For our next analysis, we examined phase–amplitude coupling (PAC) in these same regions. We focused on phase–amplitude interactions between theta and high gamma oscillations (40–70 Hz). Based upon previous observations of different patterns of PAC in the hippocampus and neocortex, we used the 2–5 Hz range in the hippocampus and 4–9 Hz in the cortex for the phase–providing frequency band.^12,34^ We compared PAC during the retrieval of contextually appropriate and inappropriate memories across subjects using the modulation index method^47^ (see Methods). Results are illustrated in Figure 4. In the posterior hippocampus and most cortical regions, we observed significantly greater PAC for correctly retrieved memories relative to PLIs (uncorrected p value < 0.01). The exception was the left inferior parietal lobule regions (AG and SMG): although it exhibited greater gamma power preceding correct recall events compared to PLIs, PAC was greater during incorrect memory retrieval (blue color in Figure 4). We extracted the preferred phase for coupling for each region during correct retrieval events, aggregated these values across subjects, and tested the phase distributions using a Rayleigh test, looking for non–uniformity in the preferred phase information.

**Figure 4:**
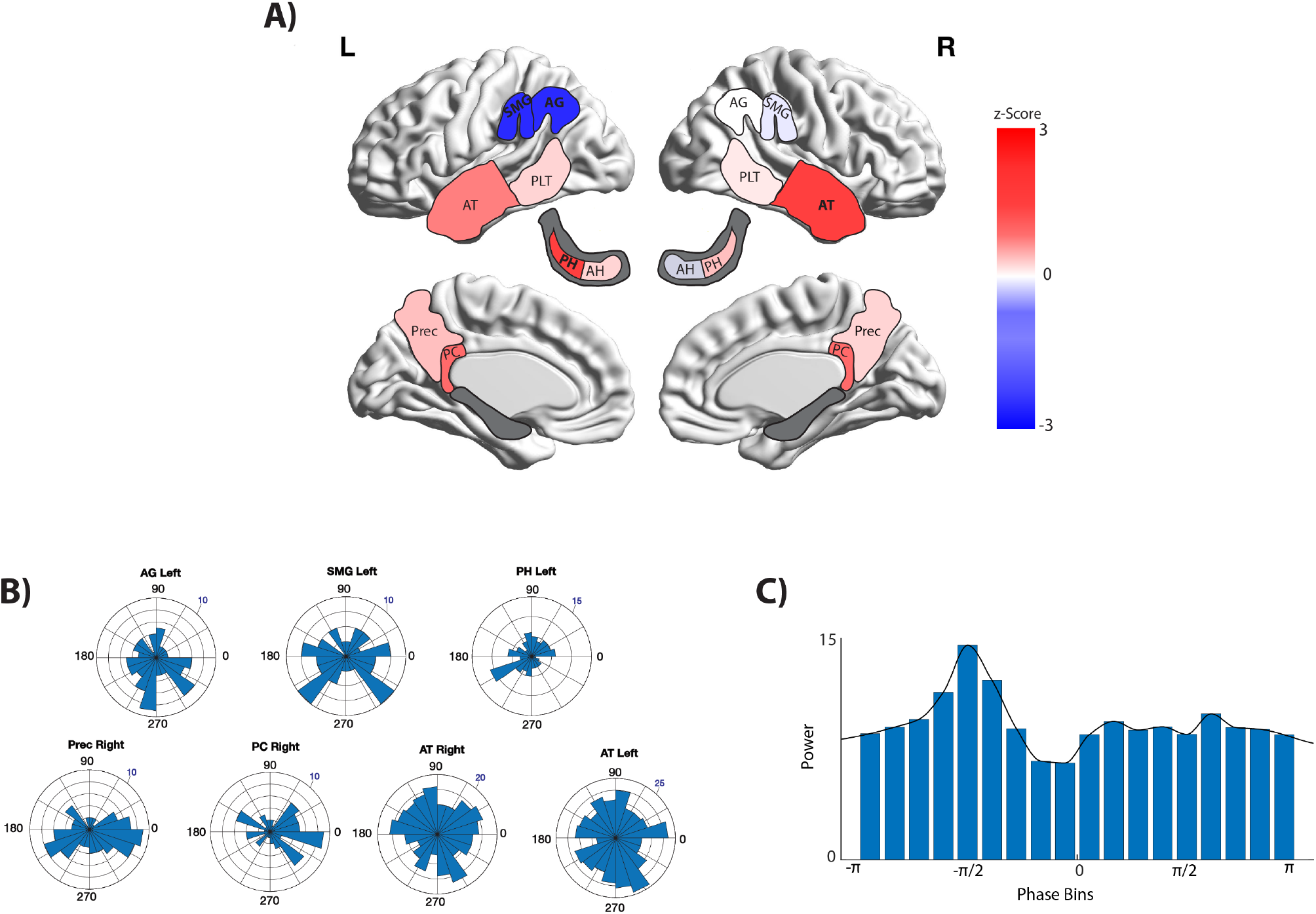
Phase–Amplitude Coupling during true versus false memory retrieval. (A) Brain plot of significant z–transformed modulation index difference between true and false memory retrieval. Warm colors indicate greater PAC during correct memory retrieval. (B) Preferred phase distribution of all significant PAC regions. Positive PAC regions (top) shows preferred phase distribution of all correctly retrieved events. Negative PAC regions (bottom) shows preferred phase distribution of PLI events. (C) Representative example of phased– binned power distribution for one trial in the posterior hippocampus (MI = 0.3).

We observed no significant effect in any region, consistent with a uniform phase distribution across subjects. Thus, although PAC distinguished between the two classes of recall events, there was heterogeneity across subjects for the preferred phase of theta–gamma coupling.

### 3.3. Phase Synchrony

Our dataset allowed us to test for connectivity relationships among our regions of interest, including cortico–hippocampal interactions. We examined connectivity in both hemispheres during correct versus incorrect retrieval, employing the PLV method to quantify connectivity differences for each subject^33,46^ in the 3–9 Hz frequency range following previous publications^46^ (see Methods). The results of these analyses are summarized in Figure 5. We observed a robust hemispheric asymmetry: connections where connectivity was elevated for correct recall vs. PLIs were more frequent in the right hemisphere (average functional connectivity 0.55 versus 1.31, *p* < 0.034) Contrary to our a priori expectations, we did not observe strong hippocampal– angular gyrus connectivity distinguishing retrieval success, although we did observe greater connectivity for the left hippocampus with the precuneus, a region that exhibited strong effects in our power analysis. The only connection for the left angular gyrus that increased with correct recall was with the prefrontal cortex (red connection in Figure 5A), a finding we address further in the *Discussion*. In the right hemisphere, angular gyrus connectivity was increased for several connections, including the precuneus, posterior cingulate, and prefrontal cortex (but not the hippocampus). The right prefrontal cortex exhibited connectivity modulated by retrieval success with two regions implicated in our analysis of HFA, namely the right angular gyrus and precuneus.

**Figure 5:**
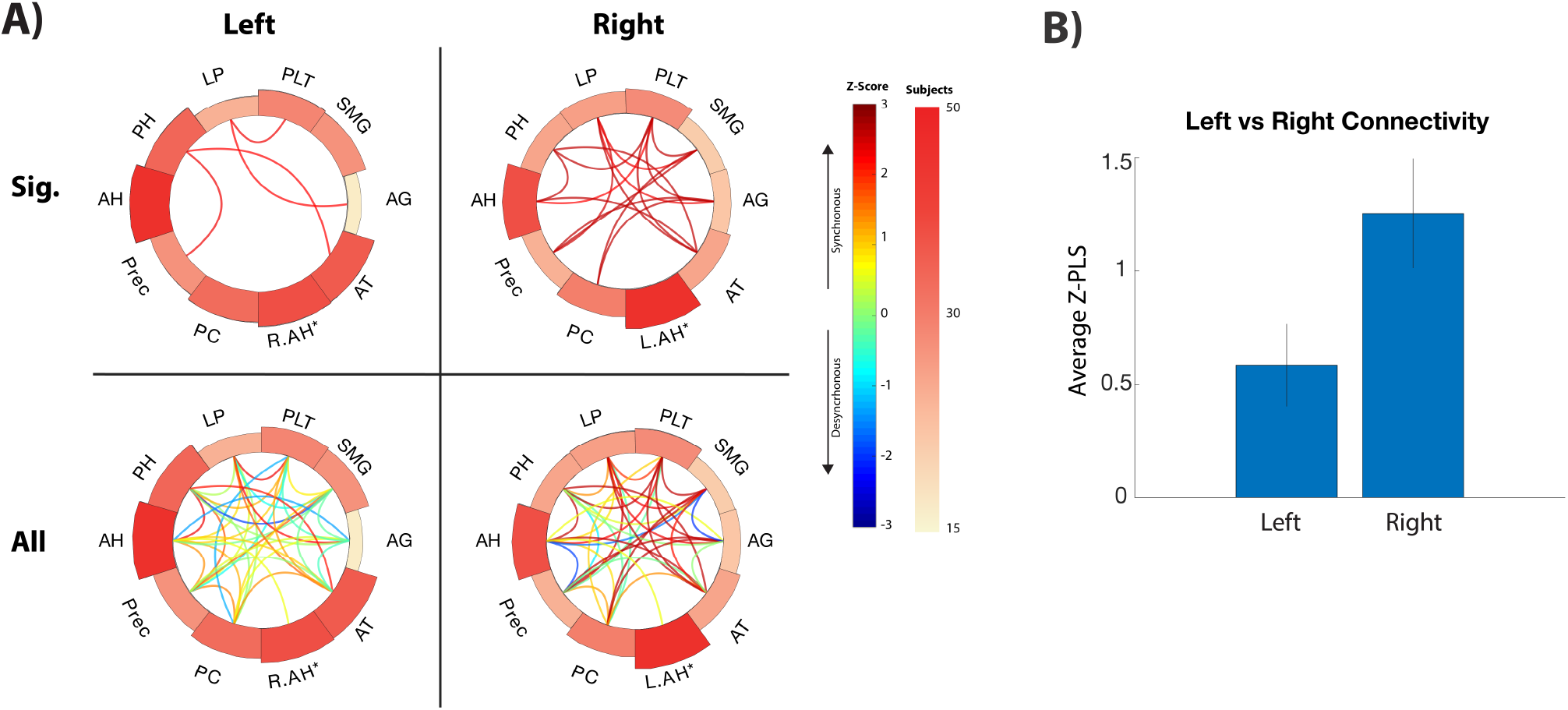
Connectivity analysis during true vs false memory retrieval. (A) Connectivity plots of retrieval network, showing connectivity difference between correct and incorrect retrieval events. Top row shows only significant connections (uncorrected *p* < 0.01, PLS difference with permutation procedure). Bottom row shows all connections regardless of magnitude. Warm colors indicate increased connectivity for successful retrieval, cool colors indicate less connectivity. Outer ring of circular plots scaled according to the number of subjects contributing data to each region. (B) Direct comparison of all left and right connectivity values aggregated across all connections. Mean PLS value shown. This was significant (*p* < 0.034).

**Figure 6:**
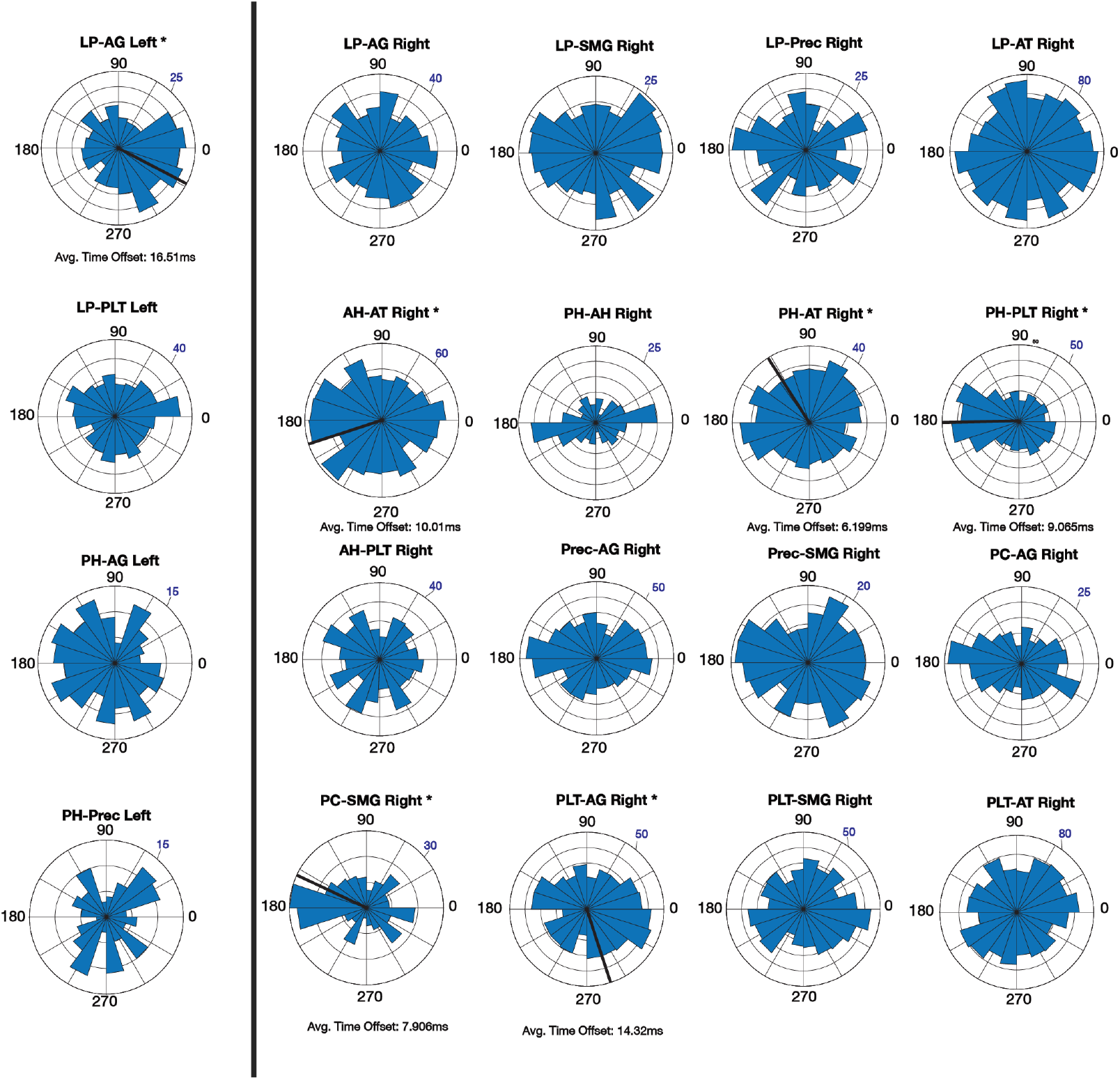
Phase Lag Distribution for significant region-region pairs. Phase lag distributions for region– region pairs. Significant non-uniform distribution of phase lags are denoted with an asterisk and provide a mean phase lag (plotted in a black line).

For region–region pairs exhibiting a significant correct versus incorrect difference in connectivity, we extracted the mean phase lag across trials (extracted at the time–point of maximum synchrony between regions for the condition of greater functional connectivity for each pair (e.g. successful retrieval), see *Methods*). The mean lag values at these timepoints were then aggregated across all electrode pairs, resulting in the rose plots illustrated in Figure 5. We observed consistent phase offsets between several functionally connected regions, most notably on the left for the angular gyrus–PFC (mean difference of 5.8 radians, uncorrected *p* < 0.001). In the right hemisphere, we observed both posterior and anterior hippocampus connections with the anterior temporal cortex exhibit a consistent phase offset (mean= 2.2 and 3.5 radians respectively, *p* < 0.05 *p* < 0.001). Furthermore, we observed that posterior hippocampus– posterior lateral temporal (mean= 3.2 Rads, *p* < 0.0001), posterior lateral temporal–angular gyrus (mean= 4.9 radians,*p* < 0.05), and posterior cingulate–supramarginal gyrus (mean = 2.7 radians, *p* < 0.001) all exhibited a strong non-uniform phase offset. Without consistent phase differences across multiple regions, we were not able to estimate the relative timing of activation across the whole retrieval network. However, we did observe that in the right temporal cortex, the phase offset for PH–posterior temporal connection was 3.2 versus 3.5 radians, which may indicate a posterior-to-anterior sequence of activation relative to the hippocampus during correct retrieval. The PLT-AG connection (4.9 radians) suggests activity in this region may follow that of the lateral temporal cortex during retrieval. We acknowledge however that interpretation of the timing of activation using these phase lags requires an assumption that connections occur within one cycle of the relevant oscillation. Characterization of timing of activation based on phase information requires a different method such as phase-slope index or dynamic causal modeling for more definitive inferences.

## 4. Discussion

fMRI investigations have provided evidence for the existence of a core episodic memory retrieval network, incorporating the hippocampus, left angular gyrus, posterior cingulate, medial prefrontal cortex, posterior middle temporal gyrus, and parahippocampal cortex.^26,50,10,19,31^ We took advantage of a unique stereo EEG data set that included a large number of intracranial electrodes in medial and lateral parietal structures, the hippocampus, and prefrontal cortex *in the same individuals* to examine oscillatory changes and functional connectivity during contextually correct versus contextually incorrect (PLIs) free recall. As compared to a previous publication^37^ (which included a subset of our data, 12/100 subjects) where only a very modest effect of recall accuracy was identified in parietal cortex, the present data set included substantially greater representation of mnemonically relevant parietal locations, including the posterior cingulate, precuneus, and angular gyrus. Overall, the pattern of HFA selectivity for correct versus incorrect memory retrieval was broadly consistent with previous reports,^37^ although the aggregation of parietal regions may have led to an attenuation of the HFA effect in previous data. However, for select regions in the parietal cortex, especially the angular gyrus and precuneus, we observed a scaling effect, by which power is greater during successful retrieval compared to both list intrusion events and activity during deliberation. By contrast, HFA in the posterior cingulate cortex was elevated both during deliberation and successful retrieval, which may be indicative of a role for this region in memory search as suggested by noninvasive data.^42^ The posterior hippocampus and angular gyrus by contrast did not exhibit this pattern.

When interpreting electrophysiological data that contrasts list intrusions with correct retrieval events in episodic memory, one must bear in mind the different scenarios that may give rise to incorrect recall. One possibility is that a study item becomes bound to the context associated with a later–occurring list,^28,52^ giving rise to a strong but contextually–inappropriate memory representation. This could occur, for example, if the study item was incidentally retrieved during the presentation of a subsequent study list, leading to false binding of the item to the new, inappropriate context. One would, however, predict that prior list intrusions generated by such false binding would elicit strong hippocampal activation, but this is inconsistent with reports that hippocampal activity discriminates between list intrusions and correctly retrieved items and is not consistent with our findings in which hippocampal activity predicts successful retrieval.^23,44,37^ The other two possibilities are 1) PLI events represent relatively weak memories that, for some reason, elude post–retrieval monitoring^21,28,52^ or 2) that PLIs represent strong, highly accessible memories that are devoid of temporal context. Our behavioral data (shown in Figure 1) suggest that PLI items are often presented (at encoding) in early serial positions. Given the link between primacy items and rehearsal, this may suggest that such ‘primacy PLI’ items represent strong but acontextual memories. This interpretation is reinforced by the tendency of some individuals to repeat PLI items across multiple lists and to retrieve a significant fraction of primacy items in the first serial position at retrieval. This tendency is consistent with the second model (strong acontextual memories), however this pattern is not universal for list intrusions: 75% of PLI events come from later serial positions during encoding and are verbalized during the latter output positions at the time of retrieval. We believe that this latter group of PLIs are likely to be more weakly encoded, consistent with model (1). It is likely that the HFA and connectivity patterns we observe reflect a mixture of these divergent mechanisms for PLI generation. Ultimately, a different episodic memory paradigm that incorporates confidence ratings may help resolve this issue.

With this in mind, we conclude from the prevalence of PLIs from later serial positions in our data (and similarity of the pattern observed in the hippocampus and AG) that the HFA recall accuracy effect in the angular gyrus constitutes direct electrophysiological evidence that AG activity supports episodic memory via participating in the representation of content of retrieved items or their context. This inference is broadly consistent with noninvasive data describing the core recollection network. The angular gyrus has been shown to exhibit BOLD signal changes that track both the amount and ‘fidelity’ of recollected information, and as such, we expected to observe successful retrieval–related effects consistent with our findings (and similar to the pattern observed in the posterior hippocampus).^16,49,51^ If by contrast the AG is more sensitive to subjective aspects of experience, we would have more likely observed similar activation for list intrusions versus successful retrieval events. The difference we identified between successful retrieval and PLI events was driven by a significant HFA decrease for list intrusions. This observation may be consistent with experiments that report increased memory errors when rTMS is applied to the angular gyrus,^45,8^ insofar as one possible effect of rTMS would be to depress HFA in this region in turn eliciting the same pattern as occurs during list intrusions in our data.

Although our findings for AG activity were broadly consistent with existing noninvasive data, it should be noted that in our data we did not observe a significant right/left difference in this effect for the angular gyrus. This comparison has not previously been tested directly to our knowledge using brain recordings. We expected that the contrast used in this experiment would elicit greater oscillatory differences for the left AG. It should be noted however that previous findings using cued recall paradigms report no clear left/right distinctions, and that studies do not often directly test left/right differences but report only those that pass statistical thresholds.^19^ Further experimentation examining left versus right angular gyrus patterns for cued retrieval paradigms may help resolve the differences in our analysis versus previous fMRI investigations of the core retrieval network, as well as to test for electrophysiological evidence that angular gyrus activity is proportional to the amount of retrieved information rather than more straightforward success–related effects.^19^

PAC in the angular gyrus has not previously been reported during memory retrieval, but our expectation was that we would observe greater PAC during correct than incorrect recall because PAC is hypothesized as a mechanism to integrate item and context information during memory retrieval.^48,20^ Insofar as PLIs represent a failure to integrate these two classes of information, lower magnitude of PAC in the angular gyrus would have been consistent with this hypothesized mechanism. This is not what we observed for the left angular gyrus, although we did identify a PAC increase during correct memory retrieval in other brain areas that exhibited an HFA recall accuracy effect (most notably, in the left posterior hippocampus). Thus, PAC may not be the mechanism by which the angular gyrus specifically integrates different features of an episodic memory representation during memory retrieval. Alternately it might be that such integration occurs on such a small spatial scale that it was undetectable given the spatial resolution afforded by our distribution of electrodes within and between participants. In relation to this point, it is noteworthy that it has been proposed that the angular gyrus comprises multiple, functionally distinct subregions.^9,51^ Finer–grained spatial sampling of angular gyrus activity, such as that afforded by 64 channel micro electrode arrays, might help resolve this question.^25^ In sum, our unexpected PAC finding for the left angular gyrus requires further investigation.

Our connectivity analysis did not uniquely identify brain regions hypothesized to participate within a core episodic retrieval network using the PLI versus correct retrieval contrast. In both hemispheres, we did not observe significant direct hippocampal–angular gyrus functional connectivity with this comparison. It should be noted that our data set did not include sufficient representation of entorhinal or parahippocampal contacts in both hemispheres to test whether these regions provide a mechanism by which parietal regions communicate with mesial temporal structures. We would expect to observe such connections, especially for the left posterior cingulate which is densely connected to the parahippocampal cortex but did not exhibit extensive connectivity in our analysis.^14^ However, we did observe an intriguing connection between the lateral prefrontal cortex and angular gyrus in both hemispheres. Given evidence of the participation of this region in the fronto–parietal control network (FCN), one intepretation of our findings is that the drop in connectivity during PLI events with the AG may be the mechanism underlying failed monitoring which then results in retrieval errors. This hypothesis will require further investigation, but is also supported by the purported role of the FCN in retrieval monitoring.^24^

The power increase we observed during PLI events in the frontal cortex has some motivation in existing noninvasive human data that have operationalized monitoring activity by eliciting weak or ambiguous memory signals.^22^ Although, in our data this power increase was visible for both correct retrieval events and list intrusions. Further clarification of prefrontal oscillatory activity during episodic memory retrieval may require paradigms that require individuals to make judgments regarding whether retrieved items are of appropriate or inappropriate context (in which individuals signal when they have retrieved an item but realize it is from the wrong list). It should be emphasized that all of our data were drawn from the lateral prefrontal cortex, as the number of midline prefrontal electrodes in our data set was relatively small and thus excluded. One might hypothesize that midline regions would be able to discriminate successfully between true and false memories based upon results from noninvasive studies,^43^ contrary to our observations. More generally, previous work suggests we should observe distinct patterns of HFA and connectivity for medial versus lateral prefrontal regions, as the former but not the latter have been implicated as part of a core retrieval network.

Perhaps our most striking observation was the hemispheric asymmetry in the connectivity data (Figure 5). We observed a greater fraction of connections significantly modulated by retrieval success in the right hemisphere, especially for connectivity increases. In itself, the finding of a hemispheric asymmetry is perhaps unsurprising given that the present free recall task relies heavily on language (although we did not observe this in our result comparing oscillatory power across conditions), but the direction of the asymmetry is arguably the opposite of what might have been anticipated. While several regions exhibited theta frequency connectivity decreases concurrent with HFA increases as previously described, this pattern was not universal in our data.^17^ HFA power and theta connectivity increases occurred simultaneously in bilateral hippocampi and the right posterior cingulate region. Additionally, an expanded data set incorporating medial PFC electrodes (or even the thalamus) may identify this area is a critical hub uniting the hippocampus (with which it has direct anatomical connectivity) and parietal lobe locations.^11^

Retrieval–related HFA was relatively greater in the posterior as compared to anterior hippocampus. A link between posterior hippocampal HFA and memory retrieval has been previously reported by us^36,35^ (using iEEG) and others^29,39^ (using fMRI data), although not specifically for correct versus incorrect recall. This finding could be interpreted as supportive of a view in which the posterior hippocampus participates in pattern completion preferentially due to greater representation of Ca3 subregions proportionally in the posterior hippocampus,^38,30^ although it should be noted that previous investigations of recollection–related effects within the retrieval network have identified activation principally in the head and body of the hippocampus.^43^ In our data set, there was not sufficient representation of electrodes localized definitively to different hippocampal subfields to test for subfield–related differences in retrieval activation.

The free recall task should provide data for neurophysiological activity during ‘bottom up’ memory search utilizing internally directed attention,^7,6^ however our data set did not include extensive electrode representation in the most dorsal aspects of the superior parietal lobule to directly compare angular gyrus versus SPL activity during deliberation. Understanding the patterns we observed and placing them within the context of a recollection network model will require further investigation, especially the comparison of free recall data with cued retrieval paradigms, as such tasks may place different demands upon internal mechanisms for cue generation preceding recollection. However, in our data, the best evidence for internal cue generation comes from the left posterior cingulate region which exhibited a significant HFA power increase during memory search (Figure 3), as well as to a lesser extent the bilateral precuneus. HFA observed in the latter structure may be consistent with the proposed contribution of the SPL as described above, although it should be noted that we observed an HFA increase preceding correct retrieval in this region as well (in fact, the largest HFA increase across all regions, Figure 3) which may imply a contribution both in memory search and the representation of features of the memory items themselves. As discussed above, these findings also relate to possible interactions among brain networks, including the core retrieval network with the frontoparietal control and salience networks. The use of brain stimulation to disrupt retrieval processes may be a useful tool for the investigation of these interactions, as suggested by findings of autobiographical^12^ and episodic memory processing. While parietal stimulation does not elicit reminiscences (unlike mesial temporal stimulation), focused disruption of specific subregions of the retrieval network (angular gyrus versus cingulate cortex versus precuneus for example) may further uncover specific contributions to retrieval processing.

## 5. Conclusion

Using a unique dataset, we demonstrate that the hippocampus, angular gyrus, and precuneus exhibit a selective HFA increase preceding contextually appropriate versus inappropriate free recall. These findings are reinforced by connectivity data, including our finding that prefrontal–parietal connectivity distinguishes correct retrieval from list intrusions. Our results inform the understanding of this important brain network and provide evidence for medial (posterior cingulate) versus lateral (angular gyrus) differences in retrieval–related activity in parietal cortex consistent with existing models describing the role of the parietal lobe in episodic memory.^6,43,13^

## References

[1] Nikolai Axmacher et al. “Interactions between medial temporal lobe, prefrontal cortex, and inferior temporal regions during visual working memory: a combined intracranial EEG and functional magnetic resonance imaging study”. In: Journal of Neuroscience 28.29 (2008), pp. 7304–7312.

[2] Jeffrey R Binder et al. “Where is the semantic system? A critical review and meta-analysis of 120 functional neuroimaging studies”. In: Cerebral cortex 19.12 (2009), pp. 2767–2796.

[3] Robert S Blumenfeld and Charan Ranganath. “Prefrontal cortex and long-term memory encoding: an integrative review of findings from neuropsychology and neuroimaging”. In: The Neuroscientist 13.3 (2007), pp. 280–291.

[4] Silvia A Bunge, B Burrows, and AD Wagner. “Prefrontal and hippocampal contributions to visual associative recognition: interactions between cognitive control and episodic retrieval”. In: Brain and cognition 56.2 (2004), pp. 141–152.

[5] John F Burke et al. “Theta and high-frequency activity mark spontaneous recall of episodic memories”. In: Journal of Neuroscience 34.34 (2014), pp. 11355–11365.

[6] Roberto Cabeza et al. “Overlapping parietal activity in memory and perception: evidence for the attention to memory model”. In: Journal of cognitive neuroscience 23.11 (2011), pp. 3209–3217.

[7] Roberto Cabeza et al. “The parietal cortex and episodic memory: an attentional account”. In: Nature Reviews Neuroscience 9.8 (2008), pp. 613–625.

[8] Paolo Capotosto et al. “Differential contribution of right and left parietal cortex to the control of spatial attention: a simultaneous EEG–rTMS study”. In: Cerebral cortex 22.2 (2012), pp. 446–454.

[9] Svenja Caspers et al. “The human inferior parietal cortex: cytoarchitectonic parcellation and interindividual variability”. In: Neuroimage 33.2 (2006), pp. 430–448.

[10] Audrey Duarte, Richard N Henson, and Kim S Graham. “Stimulus content and the neural correlates of source memory”. In: Brain research 1373 (2011), pp. 110–123.

[11] David R Euston, Aaron J Gruber, and Bruce L McNaughton. “The role of medial prefrontal cortex in memory and decision making”. In: Neuron 76.6 (2012), pp. 1057–1070.

[12] Brett L Foster and Josef Parvizi. “Resting oscillations and cross-frequency coupling in the human posteromedial cortex”. In: Neuroimage 60.1 (2012), pp. 384–391.

[13] Brett L Foster et al. “Intrinsic and task-dependent coupling of neuronal population activity in human parietal cortex”. In: Neuron 86.2 (2015), pp. 578–590.

[14] Michael F Glabus et al. “Interindividual differences in functional interactions among prefrontal, parietal and parahippocampal regions during working memory”. In: Cerebral Cortex 13.12 (2003), pp. 1352–1361.

[15] Jorge Gonzalez-Martinez et al. “Stereoelectroencephalography in the “difficult to localize” refractory focal epilepsy: early experience from a North American epilepsy center”. In: Epilepsia 54.2 (2013), pp. 323–330.

[16] Scott A Guerin and Michael B Miller. “Lateralization of the parietal old/new effect: An event-related fMRI study comparing recognition memory for words and faces”. In: Neuroimage 44.1 (2009), pp. 232–242.

[17] Simon Hanslmayr, Bernhard P Staresina, and Howard Bowman. “Oscillations and episodic memory: addressing the synchronization/desynchronization conundrum”. In: Trends in neurosciences 39.1 (2016), pp. 16–25.

[18] Hiroki R Hayama and Michael D Rugg. “Right dorsolateral prefrontal cortex is engaged during post-retrieval processing of both episodic and semantic information”. In: Neuropsychologia 47.12 (2009), pp. 2409–2416.

[19] Hiroki R Hayama, Kaia L Vilberg, and Michael D Rugg. “Overlap between the neural correlates of cued recall and source memory: evidence for a generic recollection network?” In: Journal of cognitive neuroscience 24.5 (2012), pp. 1127–1137.

[20] Melissa Hebscher, Jed A Meltzer, and Asaf Gilboa. “A causal role for the precuneus in network-wide theta and gamma oscillatory activity during complex memory retrieval”. In: ELife 8 (2019), e43114.

[21] RNA Henson, Tim Shallice, and Raymond J Dolan. “Right prefrontal cortex and episodic memory retrieval: a functional MRI test of the monitoring hypothesis”. In: Brain 122.7 (1999), pp. 1367–1381.

[22] RNA Henson et al. “Confidence in recognition memory for words: dissociating right prefrontal roles in episodic retrieval”. In: Journal of cognitive neuroscience 12.6 (2000), pp. 913–923.

[23] Marc W Howard and Michael J Kahana. “A distributed representation of temporal context”. In: Journal of Mathematical Psychology 46.3 (2002), pp. 269–299.

[24] Peggy L St Jacques, Philip A Kragel, and David C Rubin. “Dynamic neural networks supporting memory retrieval”. In: Neuroimage 57.2 (2011), pp. 608–616.

[25] Anthony I Jang et al. “Human cortical neurons in the anterior temporal lobe reinstate spiking activity during verbal memory retrieval”. In: Current Biology 27.11 (2017), pp. 1700–1705.

[26] J.D. Johnson and M.D. Rugg. “Recollection and the Reinstatement of Encoding-Related Cortical Activity”. In: Cerebral Cortex (2007).

[27] Jeffrey D Johnson, Maki Suzuki, and Michael D Rugg. “Recollection, familiarity, and content-sensitivity in lateral parietal cortex: a high-resolution fMRI study”. In: Frontiers in Human Neuroscience 7 (2013), p. 219.

[28] Marcia K Johnson. “Memory and reality.” In: American Psychologist 61.8 (2006), p. 760.

[29] Itamar Kahn et al. “Distinct cortical anatomy linked to subregions of the medial temporal lobe revealed by intrinsic functional connectivity”. In: Journal of neurophysiology 100.1 (2008), pp. 129–139.

[30] Raymond P Kesner. “Behavioral functions of the CA3 subregion of the hippocampus”. In: Learning & memory 14.11 (2007), pp. 771–781.

[31] Danielle R King et al. “Recollection-related increases in functional connectivity predict individual differences in memory accuracy”. In: Journal of Neuroscience 35.4 (2015), pp. 1763–1772.

[32] Hyunseok Kook et al. “An offline/real-time artifact rejection strategy to improve the classification of multi-channel evoked potentials”. In: Pattern Recognition 41.6 (2008), pp. 1985–1996.

[33] Jean-Philippe Lachaux et al. “Measuring phase synchrony in brain signals”. In: Human brain mapping 8.4 (1999), pp. 194–208.

[34] Bradley Lega et al. “Slow-theta-to-gamma phase–amplitude coupling in human hippocampus supports the formation of new episodic memories”. In: Cerebral Cortex 26.1 (2014), pp. 268–278.

[35] Jui-Jui Lin et al. “Gamma oscillations during episodic memory processing provide evidence for functional specialization in the longitudinal axis of the human hippocampus”. In: Hippocampus 29.2 (2019), pp. 68–72.

[36] Jui-Jui Lin et al. “Theta band power increases in the posterior hippocampus predict successful episodic memory encoding in humans”. In: Hippocampus 27.10 (2017), pp. 1040–1053.

[37] Nicole M Long et al. “Contextually mediated spontaneous retrieval is specific to the hippocampus”. In: Current Biology 27.7 (2017), pp. 1074–1079.

[38] Kazu Nakazawa et al. “Requirement for hippocampal CA3 NMDA receptors in associative memory recall”. In: Science 297.5579 (2002), pp. 211–218.

[39] Jordan Poppenk et al. “Long-axis specialization of the human hippocampus”. In: Trends in cognitive sciences 17.5 (2013), pp. 230–240.

[40] Probability and Mathematical Statistics: Statistics of Directional Data. Academic Press, London, 1972.

[41] Charan Ranganath, Marcia K Johnson, and Mark D’Esposito. “Prefrontal activity associated with working memory and episodic long-term memory”. In: Neuropsychologia 41.3 (2003), pp. 378–389.

[42] Michael D Rugg and Danielle R King. “Ventral lateral parietal cortex and episodic memory retrieval”. In: Cortex 107 (2018), pp. 238–250.

[43] Michael D Rugg and Kaia L Vilberg. “Brain networks underlying episodic memory retrieval”. In: Current opinion in neurobiology 23.2 (2013), pp. 255–260.

[44] Per B Sederberg et al. “Gamma oscillations distinguish true from false memories”. In: Psychological science 18.11 (2007), pp. 927–932.

[45] Carlo Sestieri, Gordon L Shulman, and Maurizio Corbetta. “The contribution of the human posterior parietal cortex to episodic memory”. In: Nature Reviews Neuroscience 18.3 (2017), p. 183.

[46] EA Solomon et al. “Widespread theta synchrony and high-frequency desynchronization underlies enhanced cognition”. In: Nature communications 8.1 (2017), p. 1704.

[47] Adriano BL Tort et al. “Measuring phase-amplitude coupling between neuronal oscillations of different frequencies”. In: American Journal of Physiology-Heart and Circulatory Physiology (2010).

[48] Adriano BL Tort et al. “Theta–gamma coupling increases during the learning of item– context associations”. In: Proceedings of the National Academy of Sciences 106.49 (2009), pp. 20942–20947.

[49] Kaia L Vilberg and Michael D Rugg. “Functional significance of retrieval-related activity in lateral parietal cortex: Evidence from fMRI and ERPs”. In: Human brain mapping 30.5 (2009), pp. 1490–1501.

[50] Kaia L Vilberg and Michael D Rugg. “Memory retrieval and the parietal cortex: a review of evidence from a dual-process perspective”. In: Neuropsychologia 46.7 (2008), pp. 1787–1799.

[51] Jiaojian Wang et al. “Correspondent Functional Topography of the Human Left Inferior Parietal Lobule at Rest and Under Task Revealed Using Resting-State f MRI and Coactivation Based Parcellation”. In: Human brain mapping 38.3 (2017), pp. 1659–1675.

[52] Franklin M Zaromb et al. “Temporal associations and prior-list intrusions in free recall.” In: Journal of Experimental Psychology: Learning, Memory, and Cognition 32.4 (2006), p. 792.

